# Vast amounts of encoded items nullify but do not reverse the effect of sleep on declarative memory

**DOI:** 10.1101/2020.07.23.218735

**Authors:** Luca D. Kolibius, Jan Born, Gordon B. Feld

## Abstract

Sleep strengthens memories by repeatedly reactivating associated neuron ensembles. Our studies show that although long-term memory for a medium number of word-pairs (160) benefits from sleep, a large number (320) does not. This suggest an upper limit to the amount of information that has access to sleep-dependent declarative memory consolidation, which is possibly linked to the availability of reactivation opportunities. Due to competing processes of global forgetting that are active during sleep, we hypothesised that even larger amounts of information would enhance the proportion of information that is actively forgotten during sleep. In the present study, we aimed to induce such forgetting by challenging the sleeping brain with vast amounts of to be remembered information. For this, 80 participants learned a very large number of 640 word-pairs over the course of an entire day and then either slept or stayed awake during the night. Recall was tested after another night of regular sleep. Results revealed comparable retention rates between the sleep and wake groups. Although this null-effect can be reconciled the concept of limited capacities available for sleep-dependent consolidation, it contradicts our hypothesis that sleep would increase forgetting compared to the wake group. Additional exploratory analyses relying on equivalence testing and Bayesian statistics reveal that there is evidence against sleep having a detrimental effect on the retention of declarative memory at high information loads. We argue that forgetting occurs in both wake and sleep states through different mechanisms, i.e., through increased interference and through global synaptic downscaling, respectively. Both of these processes might scale similarly with information load.

## 1 Introduction

It is undisputed that sleep is integral to the formation of long-term memory (Diekelmann & Born, 2010; Klinzing et al., 2019; Rasch & Born 2013; Walker & Stickgold, 2010). Initially, the idea prevailed that sleep predominantly acts as a passive shield against interference from novel information, as put forward by Jenkins and Dallenbach (Jenkins & Dallenbach, 1924). Even though modern interpretations of this framework still exist, it is now generally accepted that sleep plays an active role for memory (Ellenbogen, Payne, & Stickgold, 2006), with the two-stage model of memory formation (Diekelmann & Born, 2010; Marr, 1971; Klinzing, et al., 2019; Rasch & Born, 2013) being the prevailing model used in declarative memory research. First introduced by Marr (1971) it offers a solution to the “stability-plasticity-dilemma” (Abraham & Robins, 2005), which refers to the problem how a system can learn new information rapidly and in succession without overwriting older memories (Robins, 1995).

Initially the hippocampus binds together distributed information in the cortex during encoding (Battaglia et al., 2011). During subsequent sleep the hippocampus repeatedly reactivates these memories in concert with the neocortical representations (Grosmark & Buzsáki, 2016; Ji & Wilson 2007; Khodagholy et al., 2017; Ólafsdóttir et al., 2016) thereby strengthening and reorganising the representations in the neocortex (Marshall & Born, 2007; McClelland et al., 1995). Reactivation of memory traces corresponds to sharp-wave/ripple events evident in the hippocampal local field potential recordings during sleep (Diba & Buzsáki, 2007) that coordinate with sleep spindles and sleep slow oscillations to drive active systems consolidation (Clemens et al., 2007; Staresina et al., 2015; Khodagholy et al. 2017). Although, sleep spindles and reactivation in form of sharp-wave/ripples can be enhanced to compensate for large amounts of learning material (Gais et al., 2002; Mölle et al., 2009), it is plausible that an active process of sleep on memory is limited by the amount of replay that can be accommodated. This reasoning is supported by findings in humans that working memory capacity predicts declarative memory retention across sleep (Fenn & Hambrick, 2012).

In accordance with that, Feld et al. (2016) recently showed that memory consolidation of declarative content during sleep is limited in capacity. Here, participants learned either a short (40), medium (160) or long (320) list of word-pairs. Participants in the medium information load condition showed a large sleep-dependent memory advantage, whereas those in the high information load condition no longer displayed a sleep benefit. This pattern of results can be explained by a capacity limited process of active systems consolidation that leads to local potentiation of memory traces that is accompanied by a more global process of synaptic rescaling, that depotentiates synapses without being limited (Feld & Born, 2017; Tononi & Cirelli, 2014). Extrapolating from this, at even higher information loads the limited capacity for active systems consolidation is surpassed so that sleep would favour forgetting.

To test this, in the present study we doubled the information load from 320 to 640 word-pairs. We hypothesized, that sleep leads to more forgetting of word-pairs compared to a wake group.

## 2 Materials and Methods

### 3.1 Preregistration

A rough outline of this study was preregistered at AsPredicted.org. It can be viewed through this link http://aspredicted.org/blind.php?x=jc2y8t.

### 3.2 Participants

A total of 78 healthy, non-smoking, German-speaking participants performed the complete study (two participants decided to drop out prematurely). They reported a regular wake-sleep cycle, no intake of regular medication (except contraceptives) or illegal substances, and at least the qualification to enter higher education. Beginning on the morning and throughout the experiment the intake of caffeine- and alcohol-containing beverages was prohibited. Participants were randomly assigned to either the wake condition (*N* = 40; 21 female, age mean: 22.9 years, from 18-28 years) or sleep condition (*N* = 38; 19 female, age mean: 22.7 years, from 18-29 years). Participants received adequate monetary compensation for their contribution and provided written informed consent prior to the experiment. The study was approved by the local ethics committee.

### 3.3 Procedure

Participants arrived at 11:00 h and were seated in a room with four individual work-stations that were positioned to minimize distractions from other participants. See Figure 1 for a timeline of the experimental procedure. After a general instruction, participants completed a working memory capacity test (automated operation span task, OSPAN; Unsworthet al., 2005). From 12:00 to 17:00 h, participants learned the 640 word-pairs during a learning phase divided into two parts of 320 word-pairs with a short snack break in between. The snack consisted of a pretzel and one piece of fruit. Participants were asked not to actively rehearse word-pairs during breaks and oral conversation was restricted. At 17:00 h, participants received a standardized lunch consisting of either Pizza or Pasta. From 17:30 to 22:30 h, the 640 word-pairs were recalled (immediate recall) in two parts of 320 word-pairs each to estimate, how many word-pairs had been successfully encoded. Again, participants received a standardized snack in between the two parts. The snack consisted of two pieces of bread with cheese or salami and a piece of fruit. At the beginning and at the end of each experimental day as well as right before the immediate recall, the psychomotor vigilance task (PVT; Dinges et al., 1997) as well as the Stanford Sleepiness Scale (SSS; Hoddes et al., 1973) were administered (for details, see below). After recall, participants were assigned to either spend the night in the laboratory watching standardized animal documentaries (wake group) or to sleep at home (sleep group). Between 23:00 h on day 1 and 7:00 h on day 3 of the experiment, all participants wore an actigraph (ActiLife v4.4.0, ActiGraph, Pensacola, FL, USA). Participants staying in the laboratory were offered two snacks throughout the night and left the laboratory at 7:00 h. These snacks consisted of a piece of fruit, a cereal bar and two slices of raisin bread. All participants were instructed to refrain from sleeping during the day following the first experimental day. After the wake group had a recovery night, the second day of testing started at 8:00 h (delayed recall; around 33 hours after the end of the first experimental day). All participants declared compliance to the sleep-wake schedule of the experiment, which was ratified using actimetry data. At the end of the second experiment day participants had to complete a word generation task (for details, see below).

**Figure 1.**
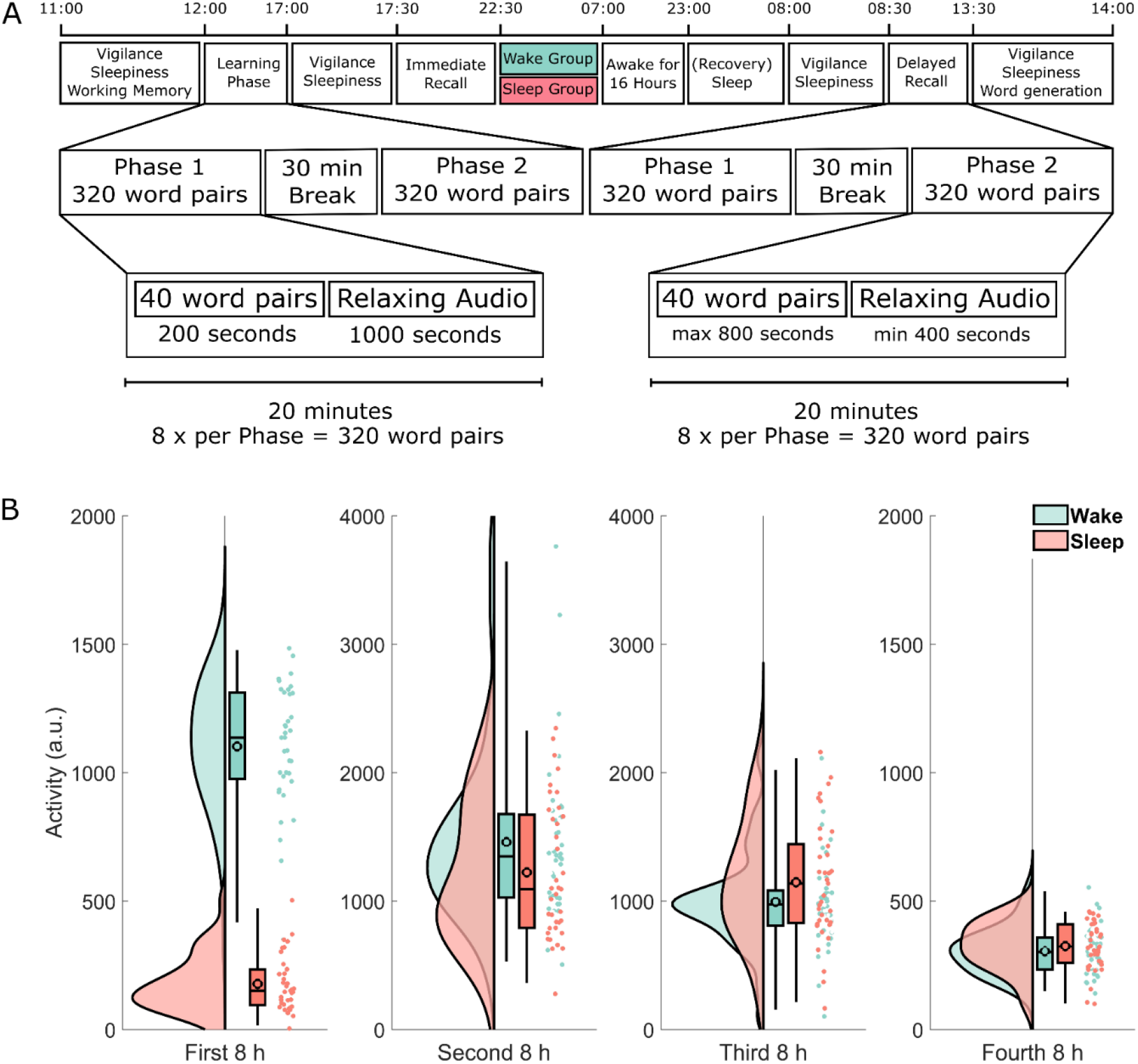
**(A)** Timeline of the experimental procedure. Learning started at 12:00 h and was followed by an immediate recall at 17:30 h as well as a delayed recall at 8:00 h two days later. The learning and recall phases each took five hours and consisted of two parts each roughly 2 hours 15 minutes long, separated by a 30 minute break. During each part, participants either learned or retrieved 320 word-pairs for a total of 640 word-pairs. Each part was further divided into eight blocks of 40 word-pairs. Each block took exactly 20 minutes. During the learning phase participants spent 3 minutes and 20 seconds per block learning word-pairs one at a time and then listened to 16 minutes and 40 seconds of relaxing audio files. During the recall phase participants had up to 20 seconds (a maximum of 13 minutes and 20 seconds per block) to respond to each of the sequentially shown cue words by typing the correct target word. Participants spent the remaining time listening to a relaxing audio file. Note that both recall phases (immediate and delayed recall) followed exactly the same procedure. **(B)** Actimetry data. Each participant was given an actigraph at the end of immediate recall to verify compliance. The y-axis shows the activity of each participant during each of the four eight-hour periods in arbitrary units. Each raincloud plot consists of the estimated distribution, a box-plot (indicating the median and the 2%, 25%, 98% quantiles, the black outlined circles depict the mean) and the activity estimations for each subject as an individual point. Data from participants in the wake group is shown in green, whereas data from the sleep group is shown in red. Note the different scale used here. See Allen et al. (2018) for the code used in this visualisation.

### 3.4 Memory task

The word-pair task was implemented using Presentation^®^ (version 16.3, Neurobehavioural Systems, Berkeley, CA, USA) on computers running on Windows 7 and adapted from Feld et al. (2016). Prior to each learning or recall block of 40 slightly related word-pairs, participants were instructed how to perform the task followed by two mock trials of the task procedure. Each phase consisted of 16 blocks which amounts to 640 word-pairs per phase. After 8 blocks there was a longer, 30-minute break. During the learning phase, each word-pair was presented once for 4s (1s inter-stimulus interval) on a horizontal axis divided by a hyphen. The left word was always the cue word, whereas the right word was always the target word. The order in which each word-pair occurred within a 40 word-pair block was randomized and the order of the word-pair blocks was balanced between participants. The block order for each participant was the same for the learning phase and both recall phases. Blocks lasted for 20 minutes. In the learning phase this meant participants were actively encoding word-pairs for 200s (40 pairs x 5s), while the remaining 1000s were spent listening to relaxing audio files. During recall, participants were presented the cue word and had 20s to type in their response. If participants wanted to move on or could not remember the target word, they were able to skip to the next word-pair by pressing the return key. Participants were instructed to answer even if they were not certain of the answer, but to avoid guesses. The keyboard input was immediately displayed which made it possible to correct for mistakes. Similar to the learning phase, recall blocks were interweaved by periods during which participants listened to relaxing audios. Again, blocks would start 20 minutes apart which resulted in a minimum of 400s of relaxing audio (if the participant used 20s to answer each cue word). The recall procedure was identical for the immediate recall after learning and the delayed recall two mornings later (Figure 1). Setting the duration of each block to 20 minutes provided two benefits: It reduced interference between the blocks and ensured an equal amount of time between each block during the learning phase and its’ corresponding block during the retrieval phase. Word-pairs were scored manually. If the answer contained spelling mistakes, used the wrong gender or number, the answer was still checked as correct. In alignment with Feld et al. (2016), we used absolute retention performance (the number of correctly recalled words at immediate recall subtracted from the number of correctly recalled words at delayed recall) as the dependent variable.

### 3.5 Working Memory (OSPAN)

The automated operation span (OSPAN; Unsworth et al., 2005) is a computer-based test to assess working memory capacity. Participants are shown simple mathematical equations in alternation with letters. Their task is to decide if the equations are correct while remembering the letters in the order they were presented. After three to six trials an array of twelve letters is shown and participants are instructed to click on the previously shown letters. For our analysis we used the absolute score and a “partial load”. The absolute score refers to the sum of all correctly recalled letters, whereas the partial load is calculated by summing up only the letters of the trials that were recalled in the correct order. One dataset was lost due to hardware malfunction. This resulted in 38 datasets in the sleep condition and 39 datasets in the wake condition.

### 3.6 Control measures (RWT, PVT, SSS, Actimetry)

In order to control for possible confounding variables, several measurements were taken. One of the detrimental effects of sleep deprivation is a diminished verbal fluency due to impaired retrieval processes (Harrison & Horne, 1997). To investigate, if, even after a recovery night, the wake condition was still suffering the negative consequences of sleep deprivation, participants completed a word generation task (RWT: Regensburger Wortflüssigkeits-Test) (Aschenbrenner et al., 2000). The assignment was to generate as many words as possible of a given category (hobbies), or to generate words that start with a specific letter (in this case “m”) during a 2-minute period. For our analysis, we added up both results for a combined score. The PVT measures the participant’s mean reaction speed and is an indicator of vigilance. The 5-minute test requires pressing the space bar, as soon as a bright red millisecond timer appears on the screen of a computer and starts counting up from 0000 in milliseconds immediately. The subject’s reaction time is displayed as soon as the space bar is pressed. For our analysis, we calculated the mean reaction speed (1 / reaction time) for each participant and the number of lapses, defined as reaction speed above 500ms (Dinges & Kribbs, 1991). Due to hard drive malfunction, we lost 6 datasets (3 in each condition) for the psychomotor vigilance task (PVT). The SSS consists of a single item measuring the subjective sleepiness on a scale of 1 (“Feeling active, vital, alert or wide awake”) to 8 (“Asleep”) (Hoddes et al., 1973). Five actimetry datasets (one from the sleep condition) could not be recovered due to hardware malfunction.

### 3.7 Statistical analysis

All statistical analysis was conducted using SPSS Version 22.0.0 on a computer running on Windows 7 and Jamovi Version 1.0.4.0. Unless stated otherwise, we relied on a univariate analysis of variance (ANOVA) with experimental condition and sex as independent variables. The SSS and the PVT were analysed using a repeated measures ANOVA with each of the five data points as a within-subjects factor and condition and sex as between-subject factors. The significance threshold for all statistical tests was set at 0.05.

## 3 Results

### 3.1 Memory Performance

Recall in the wake group decreased from *M_Wake_* = 117.85 (*SEM_Wake_* = 7.16) at immediate recall to *M_Wake_* = 106.65 (*SEM_Wake_* = 6.78) at delayed recall. In the sleep group the number of remembered word-pairs decreased from *M_Sleep_* = 112.08 (*SEM_Sleep_* = 9.93) to *M_Sleep_* = 103.61 (*SEM_Sleep_* = 9.33) at delayed recall. In a confirmatory analysis the sleep and the wake group did not differ in retention performance (the absolute amount of word-pairs that were forgotten from the immediate recall phase to the delayed recall) on the word-pair task (*M_Wake_* = −11.2 *SEM_Wake_* = 1.73, *M_Sleep_* = −8.5 *SEM_Sleep_* = 1.70; sleep/wake: *F*_1,74_ = 1.25, *p* = .27; sex: *F*_1,74_ = 0.012, *p* = 0.91; Figure 2). To provide an estimate of the evidence for the null effect we performed an equivalence test using the two one sided test procedure (Lakens et al., 2018) with upper and lower effect size bound set at *d* = ±0.2. Statistically this means we tested whether the hypotheses that the positive effect of sleep on memory retention is larger than *d* = 0.2 and that the negative effect of sleep on memory retention is larger than *d* = 0.2 can be rejected with a one sided t-tests each. Here, there was evidence that a detrimental effect of sleep on retention larger than *d* = −0.20 can be ruled out (*t*_76_ = 2.01, *p* = 0.02), whereas, there was no evidence against a positive effect of sleep larger than *d* = 0.20 (*t*_76_ = 0.24, *p* = 0.60). This means that for sleep induced forgetting there is evidence against medium and large effects, whereas small effects (*d* ≤ 0.2) could exist in our paradigm. For sleep-dependent consolidation even effects larger than *d* = 0.2 cannot be ruled in out. In addition to this analysis, following the null-hypothesis-testing (NHST) tradition, we also used Bayesian statistics to determine evidence for null effects. This analysis provides the likelihood of the model given the data, rather than the probability of the data given the model as in NHST. Similar to the equivalence test, calculating the one-sided Bayes factor provided evidence against a detrimental effect of sleep on memory, it came up with *BF*_01_ = 8.26 in favour of the null, whereas, evidence against a positive effect of sleep versus the null was undecided *BF*_01_ = 1.45. This means that the null hypothesis of no detrimental effect of sleep on memory is 8.26 times more likely than the alternative hypothesis of a detrimental effect of sleep on memory in comparison with our wake condition.

**Figure 2.**
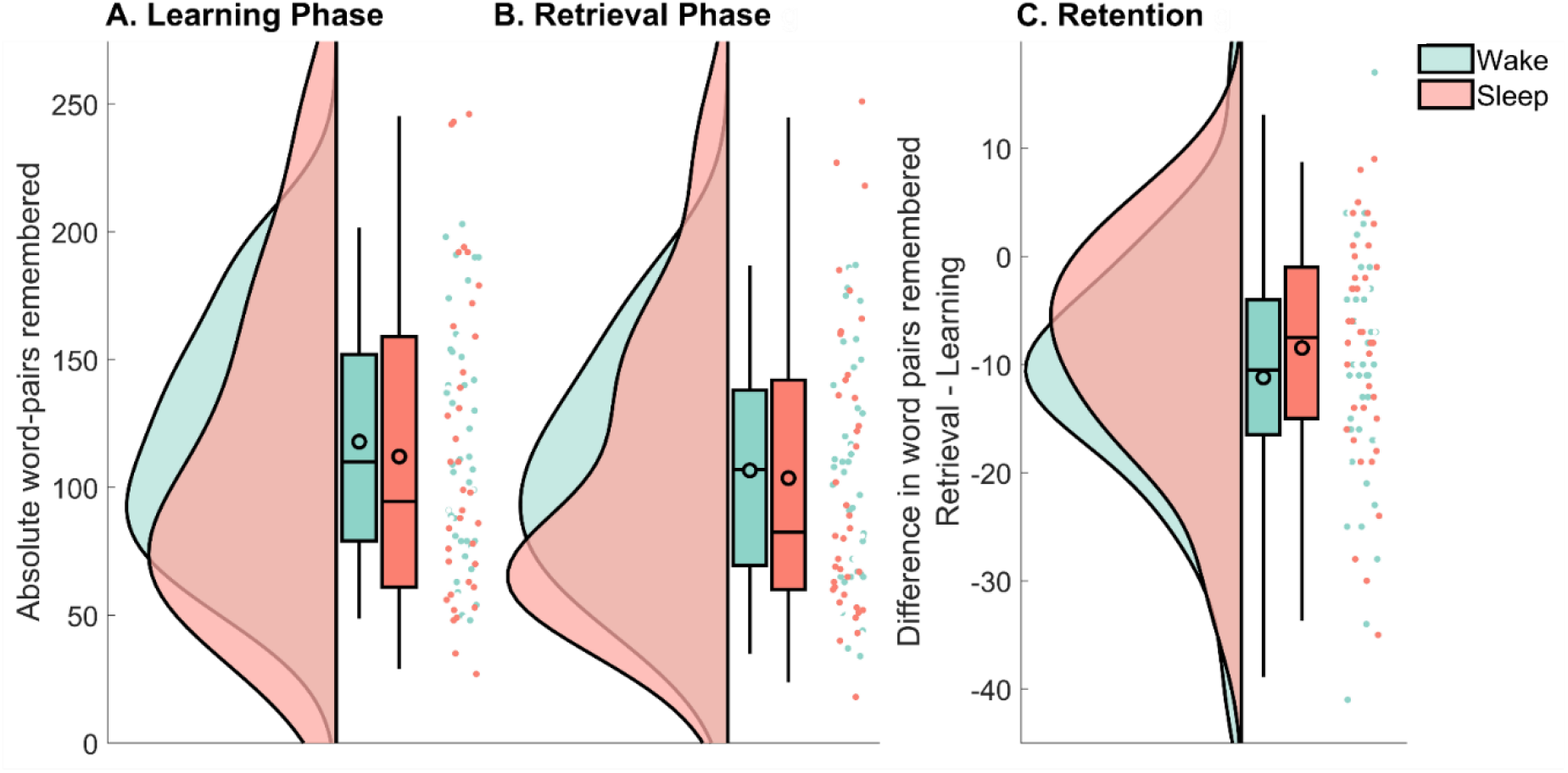
Raincloud plots (curves depict the estimated distribution, box-plots provide the median and the 2%, 25%, 75% and 98% quantiles, the black outlined circles depict the mean, the dots show individual data points) of the remembered word-pairs for the wake group (green) and the sleep group (red). **(A)** The absolute amount recalled during immediate recall that was part of the learning phase. **(B)** The absolute amount recalled during delayed recall that was part of the retrieval phase. **(C)** The difference in word-pairs remembered between retrieval and learning. Note the different scale used here. See Allen et al. (2018) for the code used in this visualisation.

Next, we investigated whether these results were due to either group having learned more words during the initial learning phase. Although the wake group descriptively learned slightly more word-pairs, this difference did not reach statistical significance (sleep/wake: *F*_1,74_ = 0.17, *p* = .68).

### 3.2 Memory Performance Blockwise

To verify this null-result, we next explored the performance per block for each condition (the experiment consisted of 16 blocks of 40 word-pairs each). As before, the absolute difference was used as a dependent variable in a repeated measures ANOVA with the block (1 to 16) as the repeated measurement and sleep/wake condition and sex as between subject variables (see Figure 3). However, no significant result emerged from this analysis (sleep/wake: *F*_1,71_ = 1.47, *p* = .22; sex: *F*_1,71_ = 0.083, *p* = .77; sleep/wake * block: *F*_15,57_ = 0.99, *p* = 0.48; sex * block: *F*_15,57_ = 0.83, *p* = 0.64).

**Figure 3.**
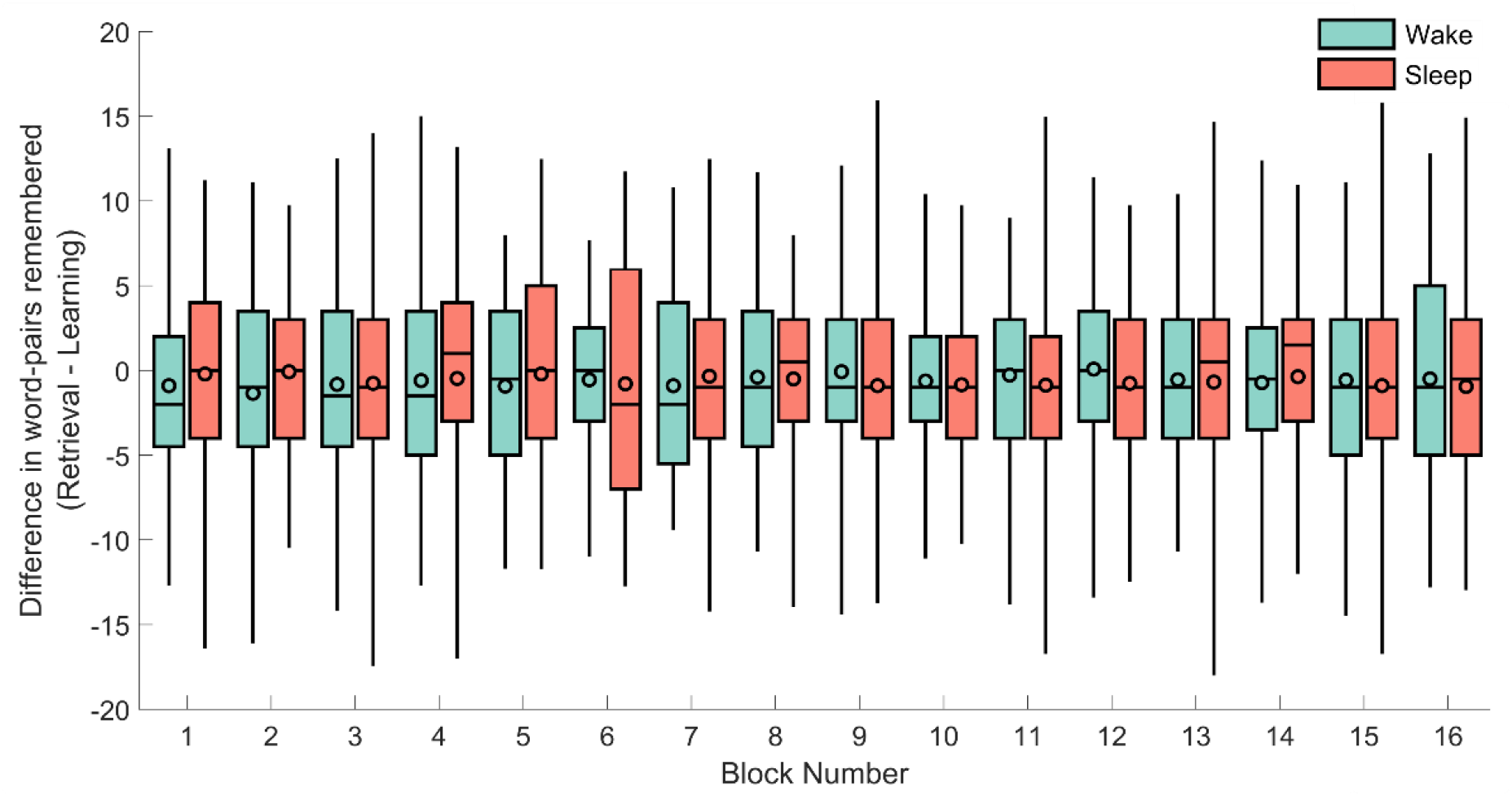
The difference in remembered word-pairs between the retrieval and learning phase for each of the individual 16 blocks for the sleep (red) and wake groups (green). The horizontal line of each box-plot indicates the median, the black outlined circles depict the mean, the border of the box indicates the 25% and 75% quartiles and the whiskers the 2% and 98% quantiles respectively.

### 3.3 Gains and Losses

Gains refer to words that were correctly recalled during delayed recall, but not during immediate recall. Losses accordingly refer to words that were correctly remembered during immediate recall, but not during delayed recall. Subjects in the sleep and wake condition both gained on average 12.3 words (*SEM_Wake_* = 0.90; *SEM_Sleep_* = 0.87). Participants in the wake group lost on average 23.5 words (*SEM_Wake_* = 1.49), while participants in the sleep group lost on average 20.79 words (*SEM_Sleep_* = 1.62). We found no statistically significant difference between groups for neither gains (sleep/wake: *F*_1,74_ = 0.001, *p* = .97), nor losses (sleep/wake: *F*_1,74_ = 1.49, *p* = .23; Table 1).

**Table 1.** Mean and SEM (in brackets) for several measures of patterns within wrong answers. Contrasted between the wake group and the sleep group.

|  | **Wake Group**<br>**Mean (SEM)** | **Sleep Group**<br>**Mean (SEM)** |
| --- | --- | --- |
| Gains<sup>1</sup> | 12.3 (0.90) | 12.3 (0.87) |
| Loss<sup>2</sup> | 23.5 (1.49) | 20.79 (1.62) |
| Number of repeated<br>wrong answers | 105.85 (6.36) | 114.03 (8.64) |
| Number of empty<br>responses<sup>3</sup> | 7.83 (14.70) | -7.42 (10.32) |
| Proportion of empty<br>responses<sup>3</sup> | 0.001 (0.026) | -0.024 (0.018) |
| Incorrect responses<sup>3</sup><br>(excluding empty<br>responses) | 3.38 (14.10) | 15.89 (9.59) |
| Word-pool error<sup>3, 4</sup> | 0.012 (0.009) | -0.006 (0.009) |

<sup>1</sup> Correctly recalled during delayed recall, but not during immediate recall
<sup>2</sup> Correctly recalled during immediate recall, but not during delayed recall
<sup>3</sup> Difference between immediate and delayed recall
<sup>4</sup> Wrong answer was learned at another point in the experiment

### 3.4 Wrong Answers

Next, we analysed the incorrect answers in detail. We investigated if there were group differences between the number of repeated wrong answers (same wrong answer during encoding and retrieval). This was not the case (*M_Sleep_* = 114.03 (*SEM_Sleep_* = 8.64), *M_Wake_* = 105.85 (*SEM_Wake_* = 6.36), sleep/wake: *F*_1,74_ = 0.64, *p* = .43; Table 1). Additionally, we explored whether the occurrence of empty responses differed between sleep/wake groups. We found no statistically significant difference between groups when considering the difference in absolute number of empty responses between immediate and delayed recall (*M_Sleep_* = −7.42 (*SEM_Sleep_* = 10.32), *M_Wake_* = 7.83 (*SEM_Wake_* = 14.70), sleep/wake: *F*_1,74_ = 0.84, *p* = .36; Table 1) or when considering the proportion of empty responses in all wrong answers (*M_Sleep_* = − 0.024 (*SEM_Sleep_* = 0.018), *M_Wake_* = 0.001 (*SEM_Wake_* = 0.026), sleep/wake: *F*_1,74_ = 0.78, *p* = .38; Table 1). Likewise, there was no significant difference between groups regarding the difference in number of incorrect responses at immediate and delayed recall. (excluding empty responses) (*M_Sleep_* = 15.89 (*SEM_Sleep_* = 9.59), *M_Wake_* = 3.38 (*SEM_Wake_* = 14.10), sleep/wake: *F*_1,74_ = 0.65, *p* = .42; Table 1).

Word-pool errors refer to wrong answers containing a word that was learned at another point in the experiment (either as a cue word or as a target word). Using the relative occurrence within all wrong answers, we calculated the difference between word-pool errors during delayed recall and during immediate recall. We found no statistically significant difference between groups (*M_Sleep_* = 0.012 (*SEM_Sleep_* = 0.009), *M_Wake_* = −0.006 (*SEM_Wake_* = 0.009), sleep/wake: *F*_1,74_ = 1.83, *p* = .18; Table 1).

### 3.5 Working Memory Capacity Task

We used a MANOVA because the two dependent variables correlated highly (*r*_75_ = 0.908, *p* < .01). The MANOVA revealed no difference in the OSPAN_Absolute_ score and the OSPAN_Partial_ score between sleep and wake condition (*M_Absolute, Sleep_* = 43.18, *SEM_Absolute, Sleep_* = 2.73; *M_Absolute, Wake_* = 46.64, *SEM_Absolut, Wake_* = 2.88; *M_Partial, Sleep_* = 60.39, *SEM_Partial, Sleep_* = 1.57; *M_Partial, Wake_* = 61.26, *SEM_Partial, Wake_* = 1.99; sleep/wake: OSPAN_Absolute_: *F*_1,73_ = 0.84, *p* = .36; OSPAN_Partial_: *F*_1,73_ = 0.12, *p* = .73; Table 2). To investigate the relation between working memory capacity and memory performance on the word-lists (using the difference score), we calculated a Pearson product-moment correlation between memory retention performance and working memory capacity separately for the sleep and wake conditions, and for the OSPAN absolute and partial score. There was a trend towards significance when looking at the wake condition and the absolute OSPAN score (*r*_39_ = −.31, *p* = .059), while all other correlations were insignificant (*p* > .11). However, when considering all participants in both groups, there was a statistically relevant negative relation between retention performance and the absolute OSPAN score (*r*_75_ = −.27, *p* = .02).

**Table 2.**
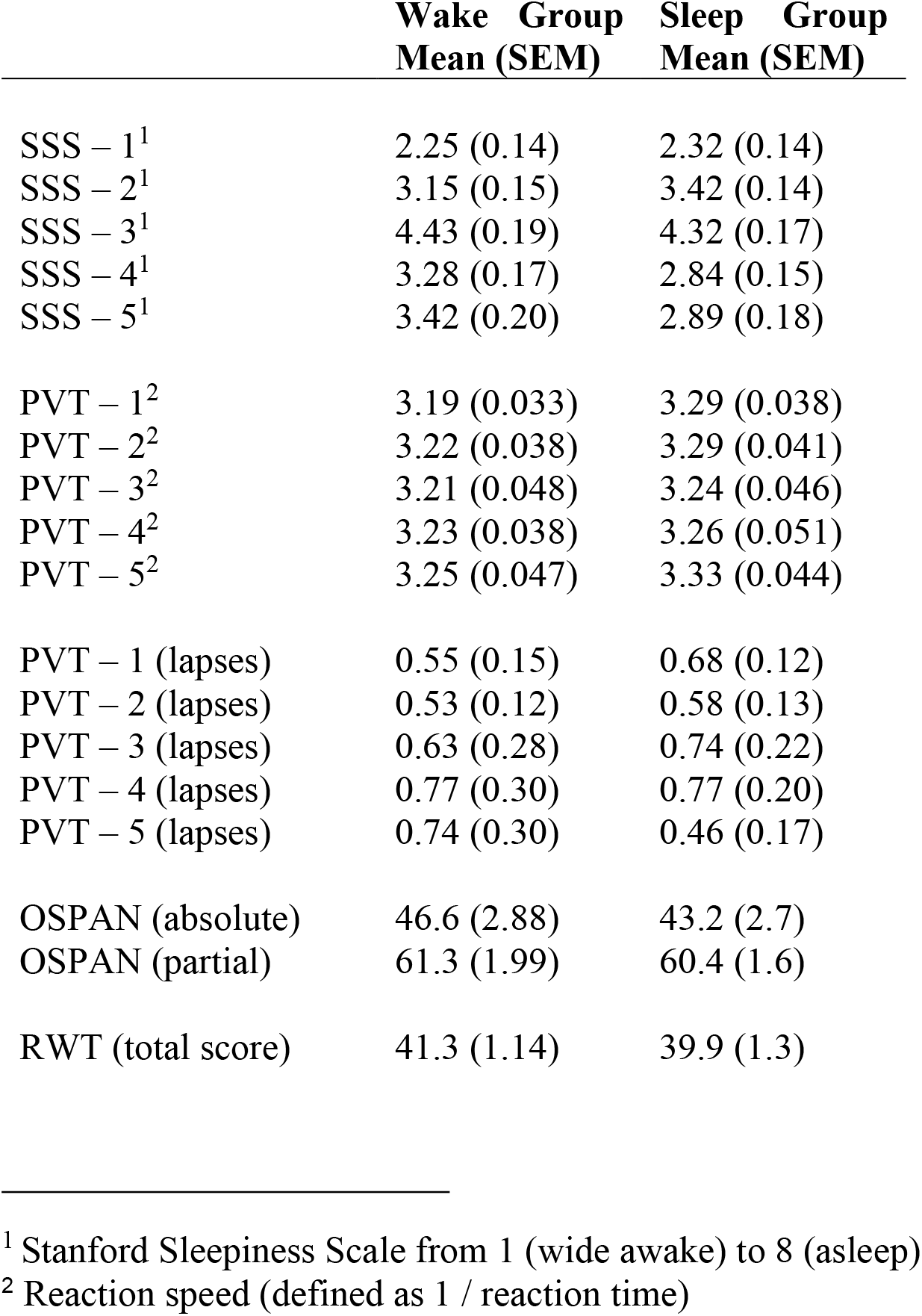
Mean and SEM (in brackets) for each of the control measures and the working memory test for the wake group and the sleep group.

|  | **Wake Group**<br>**Mean (SEM)** | **Sleep Group**<br>**Mean (SEM)** |
| --- | --- | --- |
| SSS – 1<sup>1</sup> | 2.25 (0.14) | 2.32 (0.14) |
| SSS – 2<sup>1</sup> | 3.15 (0.15) | 3.42 (0.14) |
| SSS – 3<sup>1</sup> | 4.43 (0.19) | 4.32 (0.17) |
| SSS – 4<sup>1</sup> | 3.28 (0.17) | 2.84 (0.15) |
| SSS – 5<sup>1</sup> | 3.42 (0.20) | 2.89 (0.18) |
| PVT – 1<sup>2</sup> | 3.19 (0.033) | 3.29 (0.038) |
| PVT – 2<sup>2</sup> | 3.22 (0.038) | 3.29 (0.041) |
| PVT – 3<sup>2</sup> | 3.21 (0.048) | 3.24 (0.046) |
| PVT – 4<sup>2</sup> | 3.23 (0.038) | 3.26 (0.051) |
| PVT – 5<sup>2</sup> | 3.25 (0.047) | 3.33 (0.044) |
| PVT – 1 (lapses) | 0.55 (0.15) | 0.68 (0.12) |
| PVT – 2 (lapses) | 0.53 (0.12) | 0.58 (0.13) |
| PVT – 3 (lapses) | 0.63 (0.28) | 0.74 (0.22) |
| PVT – 4 (lapses) | 0.77 (0.30) | 0.77 (0.20) |
| PVT – 5 (lapses) | 0.74 (0.30) | 0.46 (0.17) |
| OSPAN (absolute) | 46.6 (2.88) | 43.2 (2.7) |
| OSPAN (partial) | 61.3 (1.99) | 60.4 (1.6) |
| RWT (total score) | 41.3 (1.14) | 39.9 (1.3) |

<sup>1</sup> Stanford Sleepiness Scale from 1 (wide awake) to 8 (asleep)
<sup>2</sup> Reaction speed (defined as 1 / reaction time)

### 3.6 Control Measures (Actimetry, SSS, PVT, RWT)

#### 3.6.1 Actimetry Data

To analyse actimetry data, we used a repeated measures ANOVA to compare both groups’ activity levels in four 8h windows. We found a statistically significant sleep/wake * time interaction (F_1.92, 136,55_ = 35.1, p < 0.001, η^2^= .33; Greenhouse-Geisser corrected). Subsequent two-tailed t-tests revealed, that the difference was mainly driven by a difference in the first 8h window, i.e. during sleep deprivation in the wake group (*t*_48.0_ = −21.2, *p* < .001; Figure 1b).

#### 3.6.2 Subjective Sleepiness (SSS)

The SSS score at some time points was affected by the sleep/wake condition (sleep/wake * time: *F*_4,304_ = 2.46, *p* = .046, η^2^ = .031). Individual two-tailed t-tests revealed that this difference was mainly driven by higher subjective sleepiness in the wake group during the fourth (*t*_76_ = −1.92, *p* = .058) and fifth (*t*_76_ = −1.97, *p* = .053) assessment points, which both occurred on the second day of the experiment (Table 2).

#### 3.6.3 Vigilance (PVT)

There was no significant difference between groups’ average reaction speed (all *p* > .24). Likewise, there was no significant main effect or interaction of condition regarding the number of lapses (all *p* > .44; Table 2).

#### 3.6.4 Word generation task (RWT)

Analyses revealed no significant difference between conditions in the word generation task (*F*_1,74_ = 0.54, *p* = .47; Table 2).

## 4 Discussion

Previous work suggests that sleep-dependent memory consolidation is a process limited in capacity and that learning large amounts of information overloads active systems consolidation and abolishes the positive effect of sleep on memory retention (Feld et al. 2016). Extrapolating from this data, we hypothesized that at even higher loads of information during encoding sleep may favour forgetting over consolidation (Feld & Born 2017). Here, we directly tested this hypothesis by asking participants to encode a very large amount of information (640 word-pairs, twice the amount used before in the long list condition of Feld et al., 2016) before either a night of sleep or total sleep deprivation. Contrary to our predictions, we found word-pair retention to be comparable between the sleep and wake groups. While this null-effect can be reconciled with the view that capacities for consolidating memory during sleep are limited, it contradicts our hypothesis that sleep causes increased forgetting of declarative memory, when compared to wakefulness. It is important to note that sleep might still induce forgetting under conditions of massed learning, but that this effect might be masked by a direct comparison with a wake condition which itself induces forgetting (as discussed in detail below). A thorough post-hoc analysis revealed no group differences regarding a multitude of response patterns (such as gain and loss, word-pool errors and wrong answers).

In light of these results we propose that forgetting of memory traces could be achieved by different processes for the wake and sleep group, respectively. In the wake group, memory traces are more prone to interference, whereas during sleep memories are protected from interference (Ellenbogen, Hulbert et al., 2006, Wixted, 2005). However, in the sleep group, memories might be forgotten due to global synaptic downscaling (Tononi & Cirelli, 2006). In this scenario, our failure to find sleep enhanced forgetting would be explained by wake forgetting accelerating at a similar pace making it very difficult to dissociate these processes. Accordingly, if forgetting in the wake condition is primarily driven by interference within the task, then increasing or decreasing this interference (e.g. by making the stimulus material more or less semantically related) will lead to more forgetting in participants in the wake condition, but not in the sleep condition (Drosopoulos et al., 2007; Yonelinas et al., 2019). Likewise, task-unrelated interference could be manipulated by asking participants in the wake condition to learn an unrelated verbal memory task during sleep deprivation. Contrasting with the wake state, we assume that global synaptic downscaling causes forgetting in the sleep group, which could equally be manipulated in this paradigm. Since global synaptic downscaling is assumed to occur primarily during slow waves (Kim et al., 2019; Vyazovskiy et al., 2008), closed loop auditory stimulation could be employed to increase slow waves, causing more forgetting in participants that have previously encoded a high amount of information (Ngo et al., 2013). Conversely, preventing participants from reaching deep sleep should lead to less forgetting. This would prevent forgetting in two ways, first by preventing interference through new encoding, as long as sleep itself is maintained, and second by preventing synaptic downscaling during deep sleep.

An alternative account of the absence of an enhanced forgetting during sleep, induced by massed learning, can be derived by considering recent findings of synaptic downscaling mechanisms during sleep. Although it has been suggested that active systems consolidation is specific and selective, whereas synaptic downscaling is global and indiscriminate (e.g. Feld & Born, 2017), there also exists the opposite suggestion of selective downscaling during sleep (Tononi & Cirelli 2014). This latter account is largely based on findings of selective weakening of synapses during sleep, where only weaker, more plastic, synapses are erased, while stronger synapses remain stable (De Vivo et al., 2017). In addition, it has been found that sharp-wave/ripples (a correlate of memory reactivation during sleep) are involved in the depotentiation of synapses within the hippocampus (Norimoto et al., 2018), whereas they appear to be involved in the potentiation of synapses in the cortex during sleep spindles (Khodagholy et al., 2017). This offers the intriguing possibility that active systems consolidation and selective synaptic downscaling during sleep occur in a highly coordinated fashion, i.e., during the same reactivation events but in different brain areas. According to this framework the successful integration of memories into the knowledge stores of the cortex via active systems consolidation would signal the deletion of redundant memory traces from the hippocampus through synaptic downscaling. In conclusion, similar to active systems consolidation, selective synaptic downscaling during sleep might be limited by the number of available reactivations during sleep. This offers an explanation for the lack of sleep-induced net forgetting.

Turning to our findings on working memory, prior work by Fenn and Hambrick (2012) reported a positive relation between working memory and sleep related memory performance. In contrast to that, Feld et al. (2016) measured working memory performance before any sleep/wake intervention took place (which rules out any biases due to the circadian rhythms or sleep manipulation) and found no significant correlation between the two. Using the same methodology, in the present study, we found a negative correlation, although most did not reach significance. Given these contradicting findings, we suggest that sleep-dependent memory performance is likely independent of working memory functioning.

There are several limitations that were impossible to eliminate in this study and therefore possibly contribute to our results. (1) Variability in memory performance between subjects increases with list size and although the sample size was large in comparison to other studies (Ellenbogen, Hulbert et al., 2006; Igloi et al., 2015; Marshall et al., 2006; Ngo et al., 2013; Studte et al., 2017; Quigley et al., 2000) this probably decreased statistical power. To ameliorate this noise issue, criterion learning could be used in future studies, where learning is repeated until a certain percentage of word-pairs can be recalled correctly. Importantly, a study comparing different criterions found that a 60% criterion is well suited to tap into the sleep effect (Drosopoulos et al., 2007). We did not use this method, as it would have consumed significantly more time for learning, which would have made it impossible to space out learning und thus strongly increase interference effects. (2) It is possible, that not enough time had passed for sleep effects to emerge. Already Graves (1936) using nonsense-syllables found a sleep benefit only after 72h and not at shorter intervals. Similarly, Richardson and Gough (1963) did not find a sleep effect after 24h/48h, but after 144h. Especially, for large amounts of information it is conceivable that consolidation as well as forgetting is carried over to subsequent nights. (3) We tested declarative memories that were intentionally encoded. It might be that the underlying processes, such as an enhanced activation of prefrontal-hippocampal circuitry, preclude such information from sleep-dependent forgetting (Himmer et al., 2017), which stimulates the idea to compare, in future studies, sleep effects on high loads of intentionally and incidentally encoded memory.

To conclude, in the current experiment we did not find evidence that a high information load leads to more forgetting during sleep when compared to wakefulness. These findings can be explained by different mechanisms leading to forgetting in both brain states: interference-induced forgetting in the wake group and forgetting due to global synaptic downscaling in the sleep group. We propose several approaches how future studies can test this new hypothesis.

## Acknowledgments

This research was supported by grants from the Deutsche Forschungsgemeinschaft (DFG) ‘SFB 654 ‘Plasticity and Sleep’ and the European Research Council (grant No. 883098 SleepBalance) to Jan Born and a DFG Emmy Noether Grant (FE 1617/2-1) to Gordon Feld. The authors declare no competing financial interests.

## References

Abraham, W. C., & Robins, A. (2005). Memory retention–the synaptic stability versus plasticity dilemma. Trends in Neurosciences, 28(2), 73–78.

Allen, M., Poggiali, D., Whitaker, K., Marshall, T. R., & Kievit, R. (2018). Raincloud plots: a multi-platform tool for robust data visualization. PeerJ Preprints, 6, e27137v1.

Aschenbrenner, S., Tucha, O., & Lange, K. W. (2000). RWT: Regensburger Wortflüssigkeits-Test. Göttingen: Hogrefe.

Battaglia, F. P., Benchenane, K., Sirota, A., Pennartz, C. M., & Wiener, S. I. (2011). The hippocampus: hub of brain network communication for memory. Trends in Cognitive Sciences, 15(7), 310–318.

Clemens, Z., Mölle, M., Erőss, L., Barsi, P., Halász, P., & Born, J. (2007). Temporal coupling of parahippocampal ripples, sleep spindles and slow oscillations in humans. Brain, 130(11), 2868–2878.

De Vivo, L., Bellesi, M., Marshall, W., Bushong, E. A., Ellisman, M. H., Tononi, G., & Cirelli, C. (2017). Ultrastructural evidence for synaptic scaling across the wake/sleep cycle. Science, 355(6324), 507–510.

Diba, K., & Buzsáki, G. (2007). Forward and reverse hippocampal place-cell sequences during ripples. Nature Neuroscience, 10(10), 1241–1242

Diekelmann, S., & Born, J. (2010). The memory function of sleep. Nature Reviews Neuroscience, 11, 114–126. doi:10.1038/nrn2762

Dinges D., & Kribbs N. B. (1991). Performing while sleepy: effects of experimentally-induced sleepiness. In: T. H. Monk (Ed.), Human performance and cognition. Sleep, sleepiness and performance (pp. 97–128). Oxford, England: John Wiley.

Dinges, D. F., Pack, F., Williams, K., Gillen, K. A., Powell, J. W., Ott, G. E., … Pack, A. I. (1997). Cumulative sleepiness, mood disturbance, and psychomotor vigilance performance decrements during a week of sleep restricted to 4–5 hours per night. Sleep, 20(4), 267–277. doi:10.1093/sleep/20.4.267

Drosopoulos, S., Schulze, C., Fischer, S., & Born, J. (2007). Sleeps function in the spontaneous recovery and consolidation of memories. Journal of Experimental Psychology: General, 136(2), 169–183. doi:10.1037/0096-3445.136.2.169

Ellenbogen, J. M. Hulbert, J. C., Stickgold, R., Dinges, D. F., & Thompson-Schill, S. L. (2006). Interfering with theories of sleep and memory: Sleep, declarative memory, and associative interference. Current Biology, 16, 1290–1294.

Ellenbogen, J. M., Payne, J. D., & Stickgold, R. (2006). The role of sleep in declarative memory consolidation: passive, permissive, active or none?. Current Opinion in Neurobiology, 16(6), 716–722.

Feld, G. B., & Born, J. (2017). Sculpting memory during sleep: concurrent consolidation and forgetting. Current Opinion in Neurobiology, 44, 20–27. doi:10.1016/j.conb.2017.02.012

Feld, G. B., Weis, P. P., & Born, J. (2016). The limited capacity of sleep-dependent memory consolidation. Frontiers in Psychology, 7, 1–12. doi:10.3389/fpsyg.2016.01368

Fenn, K. M., & Hambrick, D. Z. (2012). Individual differences in working memory capacity predict sleep-dependent memory consolidation. Journal of Experimental Psychology: General, 141(3), 404–410. doi:10.1037/a0025268

Gais, S., Mölle, M., Helms, K., & Born, J. (2002). Learning-dependent increases in sleep spindle density. Journal of Neuroscience, 22(15), 6830–6834.

Graves, E. A. (1936). The effect of sleep on retention. Journal of Experimental Psychology, 19(3), 316–322.

Grosmark, A. D., & Buzsáki, G. (2016). Diversity in neural firing dynamics supports both rigid and learned hippocampal sequences. Science, 351(6280), 1440–1443.

Harrison, Y., & Horne, J. A. (1997). Sleep deprivation affects speech. Sleep, 20(10), 871–877.

Himmer, L., Müller, E., Gais, S., & Schönauer, M. (2017). Sleep-mediated memory consolidation depends on the level of integration at encoding. Neurobiology of Learning and Memory, 137, 101–106.

Hoddes, E., Zarcone, V., Smythe, H., Phillips, R., & Dement, W. C. (1973). Quantification of sleepiness: *A new approach*. Psychophysiology, 10(4), 431–436. doi:10.1111/j.1469-8986.1973.tb00801.x

Igloi, K., Gaggioni, G., Sterpenich, V., & Schwartz, S. (2015). A nap to recap or how reward regulates hippocampal-prefrontal memory networks during daytime sleep in humans. Elife, 4, e07903.

Jenkins, J. G., & Dallenbach, K. M. (1924). Obliviscence during sleep and waking. The American Journal of Psychology, 35(4), 605–612.

Ji, D., & Wilson, M. A. (2007). Coordinated memory replay in the visual cortex and hippocampus during sleep. Nature Neuroscience, 10(1), 100.

Khodagholy, D., Gelinas, J. N., & Buzsáki, G. (2017). Learning-enhanced coupling between ripple oscillations in association cortices and hippocampus. Science, 358(6361), 369–372.

Kim, J., Gulati, T., & Ganguly, K. (2019). Competing roles of slow oscillations and delta waves in memory consolidation versus forgetting. Cell, 179(2), 514–526.

Klinzing, J. G., Niethard, N., & Born, J. (2019). Mechanisms of systems memory consolidation during sleep. Nature Neuroscience, 22(10), 1598–1610.

Lakens, D., Scheel, A. M., & Isager, P. M. (2018). Equivalence testing for psychological research: A tutorial. Advances in Methods and Practices in Psychological Science, 1(2), 259–269.

Marr, D. (1971). Simple memory: A theory for archicortex. Philosophical Transactions of the Royal Society London B, 262, 23–81. doi:10.1098/rstb.1971.0078

Marshall, L., & Born, J. (2007). The contribution of sleep to hippocampus-dependent memory consolidation. Trends in Cognitive Sciences, 11(10), 442–450. doi:10.1016/j.tics.2007.09.001

Marshall, L., Helgadóttir, H., Mölle, M., & Born, J. (2006). Boosting slow oscillations during sleep potentiates memory. Nature, 444(7119), 610.

McClelland, J. L., McNaughton, B. L., & O’Reilly, R. C. (1995). Why there are complementary learning systems in the hippocampus and neocortex: Insights from the successes and failures of connectionist models of learning and memory. Psychological Review, 102(3), 419–457. doi:10.1037/0033-295x.102.3.419

Mölle, M., Eschenko, O., Gais, S., Sara, S. J., & Born, J. (2009). The influence of learning on sleep slow oscillations and associated spindles and ripples in humans and rats. European Journal of Neuroscience, 29(5), 1071–1081.

Ngo, H. V. V., Martinetz, T., Born, J., & Mölle, M. (2013). Auditory closed-loop stimulation of the sleep slow oscillation enhances memory. Neuron, 78(3), 545–553.

Norimoto, H., Makino, K., Gao, M., Shikano, Y., Okamoto, K., Ishikawa, T., … & Ikegaya, Y. (2018). Hippocampal ripples down-regulate synapses. Science, 359(6383), 1524–1527.

Ólafsdóttir, H. F., Carpenter, F., & Barry, C. (2016). Coordinated grid and place cell replay during rest. Nature Neuroscience, 19(6), 792.

Quigley, N., Green, J. F., Morgan, D., Idzikowski, C., & King, D. J. (2000). The effect of sleep deprivation on memory and psychomotor function in healthy volunteers. Human Psychopharmacology: Clinical and Experimental, 15(3), 171–177. doi:10.1002/(sici)1099-1077(200004)15:3<171::aid-hup155>3.0.co;2-d

Rasch, B., & Born, J. (2013). About sleep’s role in memory. Physiological Reviews, 93(2), 681–766. doi:10.1152/physrev.00032.2012

Richardson, A., & Gough, J. E. (1963). The long range effect of sleep on retention. Australian Journal of Psychology, 15(1), 37–41.

Robins, A. (1995). Catastrophic forgetting, rehearsal and pseudorehearsal. Connection Science, 7(2), 123–146.

Staresina, B. P., Bergmann, T. O., Bonnefond, M., Van Der Meij, R., Jensen, O., Deuker, L., … & Fell, J. (2015). Hierarchical nesting of slow oscillations, spindles and ripples in the human hippocampus during sleep. Nature Neuroscience, 18(11), 1679.

Studte, S., Bridger, E., & Mecklinger, A. (2017). Sleep spindles during a nap correlate with post sleep memory performance for highly rewarded word-pairs. Brain and Language, 167, 28–35.

Tononi, G., & Cirelli, C. (2006). Sleep function and synaptic homeostasis. Sleep Medicine Reviews, 10(1), 49–62. doi:10.1016/j.smrv.2005.05.002

Tononi, G., & Cirelli, C. (2014). Sleep and the price of plasticity: from synaptic and cellular homeostasis to memory consolidation and integration. Neuron, 81(1), 12–34.

Unsworth, N., Heitz, R. P., Schrock, J. C., & Engle, R. W. (2005). An automated version of the operation span task. Behavior Research Methods, 37(3), 498–505. doi:10.3758/bf03192720

Vyazovskiy, V. V., Cirelli, C., Pfister-Genskow, M., Faraguna, U., & Tononi, G. (2008). Molecular and electrophysiological evidence for net synaptic potentiation in wake and depression in sleep. Nature Neuroscience, 11(2), 200–208.

Walker, M. P., & Stickgold, R. (2010). Overnight alchemy: sleep-dependent memory evolution. Nature Reviews Neuroscience, 11(3), 218.

Wixted, J. T. (2005). A theory about why we forget what we once knew. Current Directions in Psychological Science, 14(1), 6–9.

Yonelinas, A. P., Ranganath, C., Ekstrom, A. D., & Wiltgen, B. J. (2019). A contextual binding theory of episodic memory: systems consolidation reconsidered. Nature Reviews Neuroscience, 20(6), 364–375.

